# A small molecule screen identified N-Acetyl-L-cysteine for promoting *ex vivo* growth of single circulating tumor cells

**DOI:** 10.1101/2020.08.19.257378

**Authors:** Teng Teng, Mohamed Kamal, Oihana Iriondo, Yonatan Amzaleg, Chunqiao Luo, Amal Thomas, Grace Lee, John D. Nguyen, Andrew Smith, Richard Sposto, Min Yu

**Author notes:** These authors contributed equally (Teng Teng, Mohamed Kamal).

## Abstract

Circulating tumor cells (CTCs) can be isolated via a minimally invasive blood draw and are considered a “liquid biopsy” of their originating solid tumors. CTCs contain a small subset of metastatic precursors that can form metastases in secondary organs, and provide a resource to identify mechanisms underlying metastasis-initiating properties. Despite technological advancements that allow for highly sensitive approaches of detection and isolation, CTCs are very rare and often present as single cells, posing an extreme challenge for *ex vivo* expansion after isolation. Here, using previously established patient-derived CTC lines, we performed a small molecule drug screening to identify compounds that can improve *ex vivo* culture efficiency for single CTCs. We found that N-acetylcysteine (NAC) and other antioxidants can promote *ex vivo* expansion of single CTCs, by reducing oxidative and other stress particularly at the initial stage of single cell expansion. RNA-seq analysis of growing clones and non-growing clones confirmed the effect by NAC, but also indicate that NAC-induced decrease in oxidative stress is insufficient for promoting proliferation of a subset of cells with heterogeneous quiescent and senescent features. Despite the challenge in expanding all CTCs, NAC treatment lead to establishment of single CTC clones that have similar tumorigenic features, which will facilitate future functional analyses.

## Introduction

Circulating tumor cells (CTCs) are cancer cells shed from primary or metastatic lesions into systemic circulation. Since CTCs can be shed from multiple active tumor lesions, and they contain precursors that can eventually initiate metastasis, CTCs are considered a liquid biopsy for solid tumors^1,2^. It has been shown that high numbers of CTCs correlate with a worse prognosis in several types of cancer^1,3^. Despite significant variability between patients and disease stages, CTCs are generally very rare. Most patients with metastatic cancers, including prostate, ovarian, breast, gastric, colorectal, bladder, renal, non-small cell lung, and pancreatic cancers, have low numbers of CTCs in a tube of blood, according to an analysis using the CellSearch platform, which captures CTCs based on EpCAM expression^4^. Although technologies that do not solely rely on EpCAM-surface expression have been reported to capture a higher number of CTCs^1,5,6^, the quantity remains too low for downstream functional analysis in most cases.

Several studies have shown successful *ex vivo* expansion of CTCs isolated from patients with breast^7^, colorectal^8^, and prostate^9^ cancer. These CTC lines have provided sufficient amounts of material for many analyses, including xenograft analysis and drug susceptibility assessment^7,10^. However, the efficiency of establishing the *ex vivo* culture of CTCs is extremely low, limiting its broad application to the majority of the cancer patients^11^. This low efficiency may be due to limited quantities, low capture efficiency, the harshness of the procedure, and the vulnerability of CTCs in circulation^11^. It has been shown that CTCs experience significant stress from high reactive oxygen species (ROS) levels, induced by the detachment of the extracellular matrix (ECM) or cell-cell connections^12-16^. Cells resilient to ROS may have a higher chance of initiating metastasis^12,13,16-19^. Changes in both glucose and glutamine metabolism have been found to regulate proper redox balance in CTCs and promote anchorage independent growth^20^. Antioxidants have recently been shown to promote CTC survival and metastasis in lung, melanoma, breast, and prostate cancers^12,13,16-19^. In addition, CTC clusters have a greater tendency to survive and metastasize, due to cell-cell interactions^21-23^. However, CTC clusters are only detected in a small percentage of patients, while single CTCs are far more common. Improved culture conditions for expanding single CTCs may help resolve their molecular and phenotypic properties.

Due to the rarity of CTCs, optimizing culture conditions for single cell expansion has been challenging. We previously reported a method for the *ex vivo* culture of CTCs derived from metastatic breast cancer patients and established several CTC lines^7^. These CTC lines exhibit metastatic potential that represents the major metastatic lesions in corresponding patients^10^. In addition, they are cultured in suspension and physiological oxygen level, mimicking the environment of CTCs in venous blood circulation. Although these CTC lines can be maintained long term in culture, when dissociated and plated as single cells, it is extremely difficult for majority of these cells to expand successfully. Hence, we used single CTCs as a platform to screen a small molecule library to identify compounds that promote their expansion in culture.

## Results

### Initial low-confidence screening for single CTC expansion

We first performed a low-confidence initial screen using single CTCs sorted from our CTC lines with 317 compounds from the StemSelect library (Supplementary Table S1). We sorted single live CTC into each well of a 96-well plate. The wells containing single cells were confirmed under a light microscope on the day of sorting and used for the screen. Cell numbers in these wells were quantified every 6 days, and media containing small molecules was replenished every 3 days (Figure 1). This initial screen was performed with 12-18 wells per compound, totaling 28 successful batches with 235 plates from 3 patient-derived CTC lines (BRx68, BRx07, and BRx50). DMSO controls were included in each batch and the compounds were tested blindly with code names. Due to significant heterogeneity in single cells, the limited number of wells tested for each compound in this initial screen is not sufficient for statistical analysis. Therefore, to identify a recurring pattern of compounds that function in similar pathways, we analyzed all the compounds that showed a higher median value of growth, based on area under the curve (AUC) calculation of cell numbers over time, compared to controls in the same batch, including CTC media only (CT) or media with vehicle DMSO (CD). Among the 130 compounds detected to have a higher median AUC than the controls (Table 1, Supplementary Table S2, S3), we found many compounds with similar biological functions. Seven out of 12 cyclooxygenase (COX) inhibitors, 6 out of 11 antioxidants and free radical scavengers, and 4 out of 4 5’ adenosine monophosphate-activated protein kinase (AMPK) activators increased CTC growth in this initial screen (Table 1). Since all 3 pathways have been previously linked with reducing cellular ROS levels^13,18,24^ and given recent reports showing the role of antioxidants in CTC survival, we also tested a commonly used antioxidant N-Acetyl-L-cysteine (NAC). We first evaluated the effect of several different concentrations of NAC on the BRx68 line in two different batches and found the 200 μM-300 μM NAC showed the most significant effect in promoting single CTC proliferation (Supplementary Figure 1). Therefore, we selected NAC and several different compounds from these pathways to further validate in a second-round test using a larger number of wells for statistical evaluation.

**Table 1.**
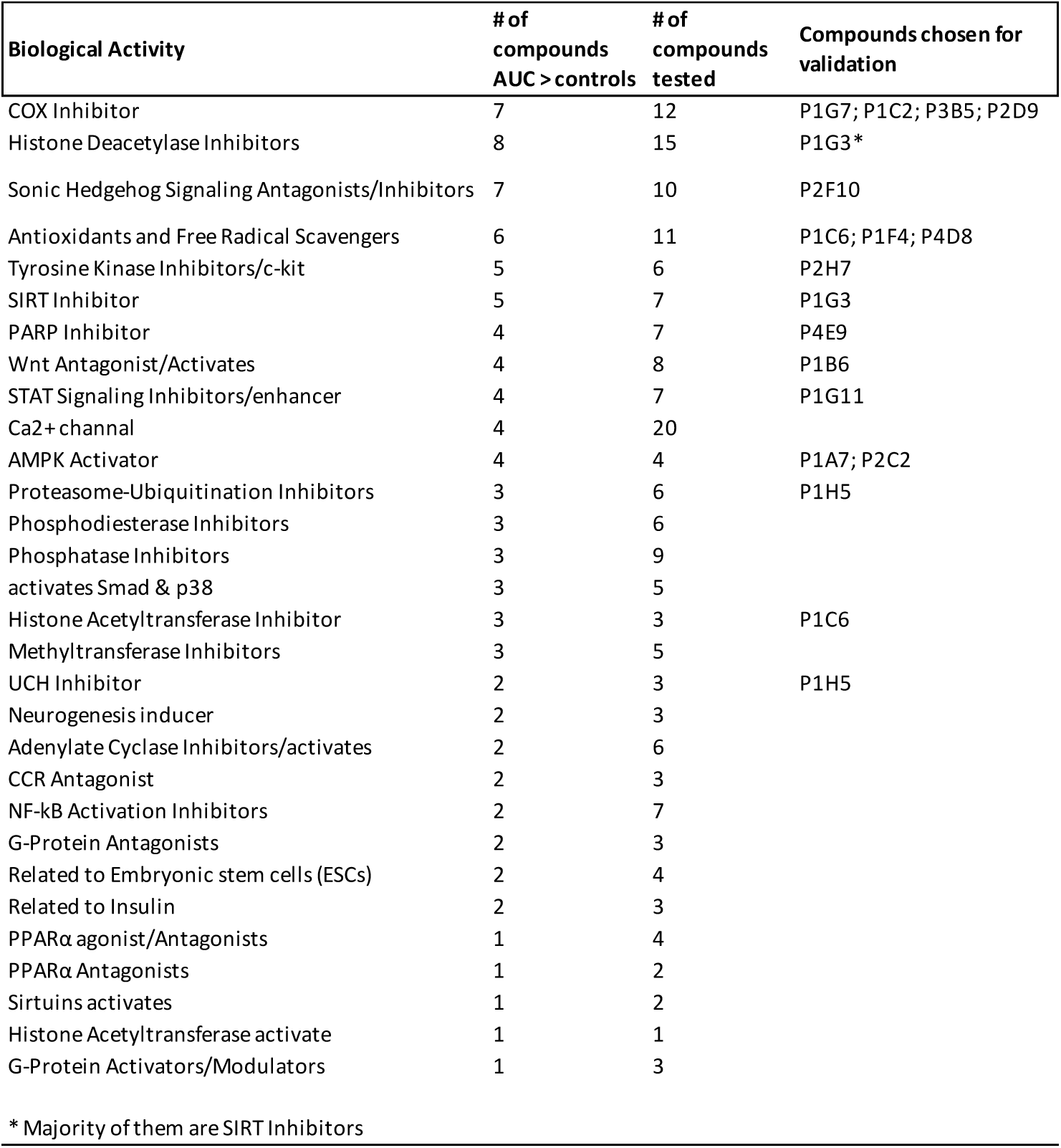
Analysis of a list of identified compounds from the initial screen.

| **Biological Activity** | **# of compounds AUC > controls** | **# of compounds tested** | **Compounds chosen for validation** |
| --- | --- | --- | --- |
| COX Inhibitor | 7 | 12 | P1G7; P1C2; P3B5; P2D9 |
| Histone Deacetylase Inhibitors | 8 | 15 | P1G3* |
| Sonic Hedgehog Signaling Antagonists/Inhibitors | 7 | 10 | P2F10 |
| Antioxidants and Free Radical Scavengers | 6 | 11 | P1C6; P1F4; P4D8 |
| Tyrosine Kinase Inhibitors/c-kit | 5 | 6 | P2H7 |
| SIRT Inhibitor | 5 | 7 | P1G3 |
| PARP Inhibitor | 4 | 7 | P4E9 |
| Wnt Antagonist/Activates | 4 | 8 | P1B6 |
| STAT Signaling Inhibitors/enhancer | 4 | 7 | P1G11 |
| Ca<sup>2+</sup> channel | 4 | 20 |  |
| AMPK Activator | 4 | 4 | P1A7; P2C2 |
| Proteasome-Ubiquitination Inhibitors | 3 | 6 | P1H5 |
| Phosphodiesterase Inhibitors | 3 | 6 |  |
| Phosphatase Inhibitors | 3 | 9 |  |
| activates Smad & p38 | 3 | 5 |  |
| Histone Acetyltransferase Inhibitor | 3 | 3 | P1C6 |
| Methyltransferase Inhibitors | 3 | 5 |  |
| UCH Inhibitor | 2 | 3 | P1H5 |
| Neurogenesis inducer | 2 | 3 |  |
| Adenylate Cyclase Inhibitors/activates | 2 | 6 |  |
| CCR Antagonist | 2 | 3 |  |
| NF-κB Activation Inhibitors | 2 | 7 | |
| G-Protein Antagonists | 2 | 3 |  |
| Related to Embryonic stem cells (ESCs) | 2 | 4 |  |
| Related to Insulin | 2 | 3 |  |
| PPARα agonist/Antagonists | 1 | 4 | |
| PPARα Antagonists | 1 | 2 | |
| Sirtuins activates | 1 | 2 |  |
| Histone Acetyltransferase activate | 1 | 1 |  |
| G-Protein Activators/Modulators | 1 | 3 |  |

\* Majority of them are SIRT Inhibitors

**Figure 1.**
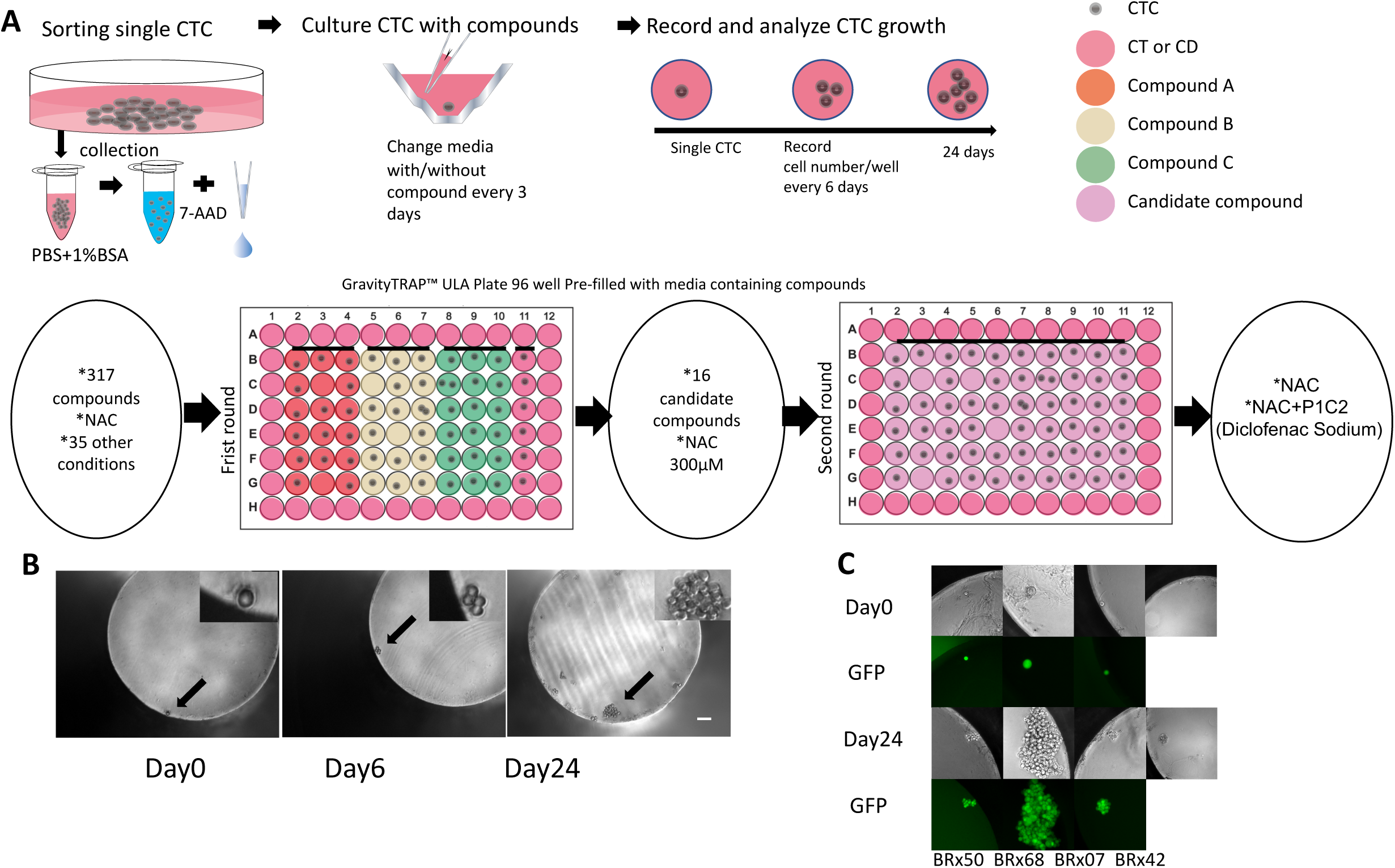
Illustration of the single cell drug screen process. **(A)** Illustration of the small molecule screening process (top panel) and the summary of results from the first and second round screenings (bottom panel). **(B)** Representative phase contrast images of the growth of a single BRx68 CTC at different time points. Scale bar: 200μm. **(C)** Representative phase contrast and GFP-fluorescent images of CTC clones generated from different CTC lines (BRx50, BRx68, BRx07, and BRx42). BRx42 cells used are not GFP-transduced.

### Validation of candidate compounds

We selected 16 compounds plus NAC from the most promising biological activity categories to validate in 4 patient-derived CTC lines (BRx07, BRx42, BRx50, and BRx68), using 36 different concentrations or combinations. This totaled 13 batches consisting 166 plates with 60 wells per plate. Similar to the initial screen, media were changed every 3 days, and cell numbers in each well were counted every 6 days until day 24.

Results showed that the best compounds that are universal to all 4 CTC lines are NAC at 300μM, or NAC (300μM) in combination with the P1C2 compound—a COX-1/2 inhibitor, Diclofenac Sodium (1μM or 0.5μM) (Table 2, Supplementary Table S4, S5). Compared to controls, NAC or NAC+P1C2 consistently showed statistically significant improvement for single CTC expansion across many different batches for all 4 CTC lines (Figure 2A&B, Table 2). For BRx50 and BRx42 lines, which are extremely difficult to expand as single cells, the addition of these compounds can lead to the successful generation of single cell clones. Moreover, we identified compounds that increased CTC expansion in a cell line-specific manner. For example, the P1G3 compound (AGK2, a reversible inhibitor for Sirtuin-2 (SIRT2), a subclass of histone deacetylase inhibitors) can promote single cell growth specifically in BRx42 (Figure 2C), while the P4D8 compound (LY 231617, an antioxidant and free radical scavenger) can promote single cell growth for the BRx68 line, but inhibit growth for BRx07 and show a tendency towards decreased growth in other lines (Figure 2D, Supplementary Table 4).

**Table 2.** Results of a list of conditions validated in the second-round screen.

| Compound | Biological Activity | Concentration (μM) | Batch | Patient | Control | Drug (N) | Control (N) | Drug Median AUC | Control Median AUC | P | Adjusted P | Note |
| --- | --- | --- | --- | --- | --- | --- | --- | --- | --- | --- | --- | --- |
| NAC | Antioxidants and Free Radical Scavengers | 300 | 12/21/17 | BRx50 | CT | 51 | 96 | 138 | 30 | <0.001 | <0.001 |  |
| NAC |  | 300 | 9/26/18 | BRx50 | CD | 43 | 46 | 3 | 0 | <0.001 | 0.0019 |  |
| NAC |  | 300 | 5/17/18 | BRx50 | CD | 45 | 38 | 120 | 31.5 | <0.001 | 0.0046 |  |
| NAC |  | 300 | 4/19/18 | BRx50 | CD | 47 | 41 | 123 | 36 | 0.0029 | 0.0224 |  |
| NAC |  | 300 | 5/17/18 | BRx50 | CT | 45 | 44 | 120 | 33 | 0.0031 | 0.0227 |  |
| NAC |  | 300 | 4/19/18 | BRx42 | CT | 46 | 41 | 21 | 9 | 0.0041 | 0.0289 |  |
| NAC |  | 300 | 4/17/18 | BRx68 | CT | 47 | 76 | 63 | 21 | 0.0187 | 0.0864 |  |
| NAC |  | 300 | 11/16/17 | BRx68 | CT | 44 | 88 | 52.5 | 31.5 | 0.0486 | 0.1743 |  |
| NAC |  | 300 | 4/19/18 | BRx42 | CD | 46 | 51 | 21 | 9 | 0.0515 | 0.1743 |  |
| NAC |  | 300 | 2/13/18 | BRx07 | CT | 49 | 103 | 90 | 48 | 0.061 | 0.1952 |  |
| NAC |  | 300 | 9/26/18 | BRx68 | CT | 40 | 45 | 130.5 | 24 | 0.0631 | 0.1952 |  |
| NAC |  | 300 | 9/26/18 | BRx68 | CD | 40 | 47 | 130.5 | 18 | 0.0651 | 0.1952 |  |
| NAC |  | 300 | 4/17/18 | BRx07 | CT | 47 | 77 | 78 | 42 | 0.0652 | 0.1952 |  |
| NAC |  | 250 | 12/14/17 | BRx68 | CT | 47 | 99 | 81 | 33 | 0.0666 | 0.1955 |  |
| NAC+P1C2 | Antioxidants and Free Radical Scavengers + COX Inhibitor | 300 + 1 | 2/15/18 | BRx42 | CT | 45 | 88 | 9 | 0 | <0.001 | <0.001 |  |
| NAC+P1C2 |  | 300 + 1 | 4/19/18 | BRx50 | CD | 45 | 41 | 141 | 36 | <0.001 | 0.0022 |  |
| NAC+P1C2 |  | 300 + 1 | 4/19/18 | BRx50 | CT | 45 | 49 | 141 | 66 | 0.0306 | 0.1222 |  |
| NAC+P1C2 |  | 300 + 1 | 2/13/18 | BRx07 | CT | 53 | 103 | 72 | 48 | 0.0525 | 0.1743 |  |
| NAC+P1C2 |  | 300 + 0.5 | 5/17/18 | BRx07 | CT | 46 | 48 | 123 | 51 | <0.001 | 0.0031 |  |
| NAC+P1C2 |  | 300 + 0.5 | 5/17/18 | BRx07 | CD | 46 | 51 | 123 | 48 | 0.0029 | 0.0224 |  |
| NAC+P1C2 |  | 300 + 0.5 | 5/17/18 | BRx50 | CD | 44 | 38 | 132 | 31.5 | 0.005 | 0.0325 |  |
| NAC+P1C2 |  | 300 + 0.5 | 5/17/18 | BRx50 | CT | 44 | 44 | 132 | 33 | 0.0195 | 0.0873 |  |
| NAC+P1C2 |  | 300 + 0.5 | 5/17/18 | BRx68 | CT | 43 | 45 | 678 | 342 | 0.0198 | 0.0873 |  |
| NAC+P1C2 |  | 300 + 0.5 | 5/17/18 | BRx68 | CD | 43 | 45 | 678 | 402 | 0.0517 | 0.1743 |  |
| P1C2 | COX Inhibitor | 1 | 4/19/18 | BRx42 | CT | 49 | 41 | 18 | 9 | 0.0099 | 0.0526 |  |
| P1C2 |  | 1 | 5/17/18 | BRx50 | CD | 40 | 38 | 108 | 31.5 | 0.0103 | 0.0533 |  |
| P1C2 |  | 1 | 5/17/18 | BRx50 | CT | 40 | 44 | 108 | 33 | 0.0362 | 0.1385 |  |
| P1G3 | SIRT Inhibitor | 1 | 2/15/18 | BRx42 | CT | 46 | 88 | 6 | 0 | 0.0047 | 0.0321 |  |
| P1G3 |  | 1 | 4/19/18 | BRx42 | CT | 48 | 41 | 15 | 9 | 0.0063 | 0.0344 |  |
| P3B5 | COX Inhibitor | 1 | 9/26/18 | BRx50 | CD | 28 | 46 | 6 | 0 | <0.001 | <0.001 | * |
| P4D8 | Antioxidants and Free<br>Radical Scavengers | 1 | 11/16/17 | BRx68 | CT | 53 | 88 | 111 | 31.5 | 0.0053 | 0.0332 |  |

\* no more drug available on the mark

**Figure 2.**
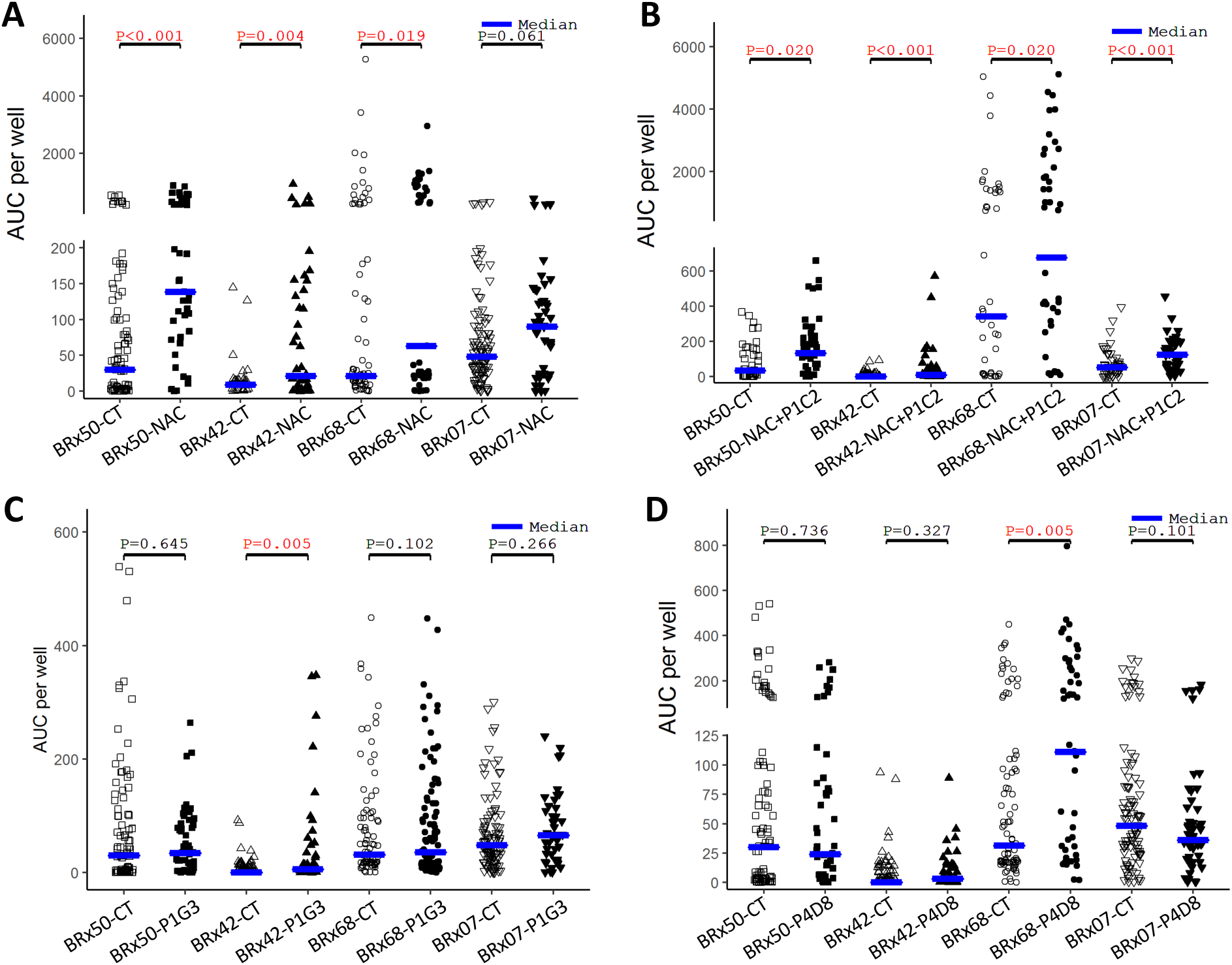
NAC and NAC+P1C2 combined promote growth of single CTCs from multiple lines. Graph showing AUC measurement of the proliferation of single CTCs from 4 different CTC lines over 24 days with NAC 300 μM **(A)**, NAC + P1C2 **(B**), P1G3 **(C)**, or P4D8 **(D)**. * *P*<0.05; ** *P*<0.01; *** *P*<0.001. *P* values were obtained by a Kruskal–Wallis test adjusted by Benjamini-Hochberg Procedure for multiple testing.

To mimic the CTC isolation procedure, we performed spiked-in experiments with 3 CTC lines (BRx42, BRx50 and BRx68) into healthy donors’ blood. We used the RosettSep CTC isolation method to isolate the spiked CTCs, and separated isolated CTCs equally into wells with control media or with NAC or NAC plus P1C2. Compared to the control condition, NAC or NAC plus P1C2 significantly increased the growth of isolated CTCs (Supplementary Figure 2). This confirmed the effects of these compounds on expanding CTCs that have been processed through an isolation procedure.

### NAC single cell clones form tumors with similar kinetics as controls

We noticed that the most critical phase is the initial expansion from single CTCs. Once a single cell colony reaches a critical size, treatment with these compounds (NAC or NAC+P1C2) does not seem to confer additional growth advantages. Therefore, we stopped the treatment of these compounds at 24 days, and then prolonged the culture to generate several single cell clones. To evaluate the tumorigenicity of the single cell clones, we injected GFP-Luciferase tagged BRx68 single clones generated with NAC treatment or control clones into the mammary fat pads of female NSG mice. The NAC treated clones generated tumors with similar growth kinetics and histology, indicating that short term treatment with NAC did not significantly affect CTCs’ tumorigenicity (Figure 3).

**Figure 3.**
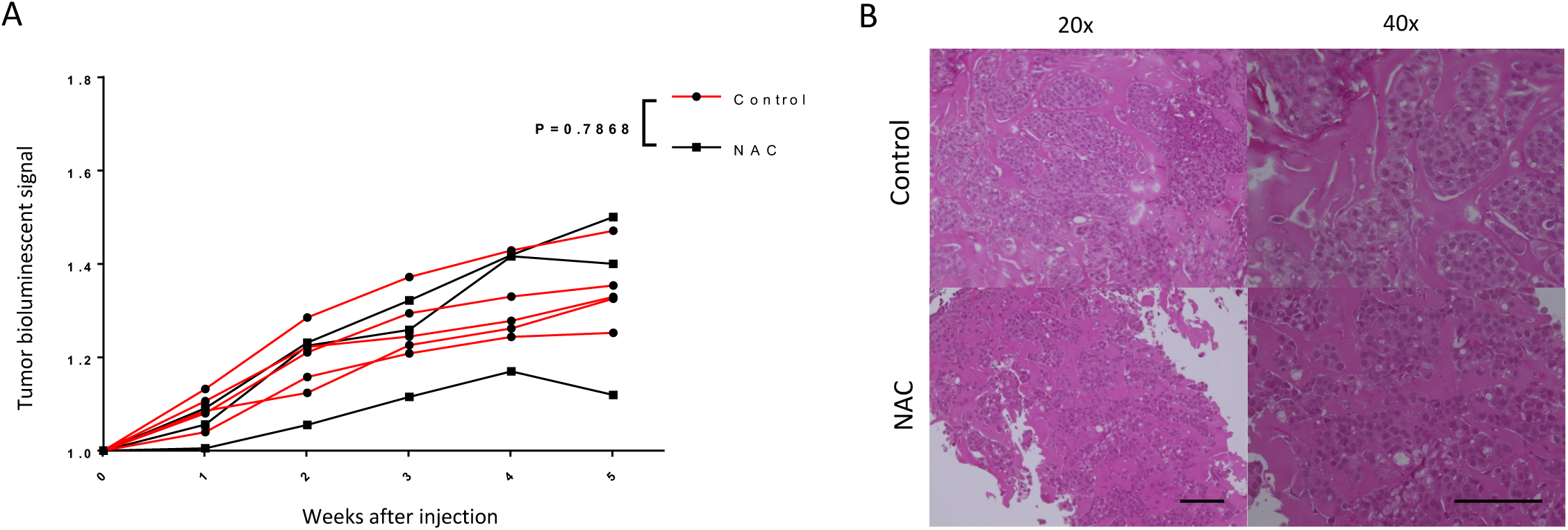
Pretreatment with short time NAC dose not change the tumorigenicity of CTCs. **(A)** The graph shows the tumor growth kinetics of single BRx68 clones generated with (NAC) or without (control) NAC after the first 24 days (NAC group: N=3, control group N=5). *P*=0.7868. *P* value was analyzed by two-way ANOVA with RM by columns between 2 groups at matched time point. Interaction between groups has been tested. **(B)** Representative images of Hematoxylin & Eosin staining of the primary tumor generated in control and NAC groups. Scale bar: 100 μm.

### NAC rescued the proliferation of freshly isolated CTCs from a breast cancer patient

We used our newly developed negative selection method of PIC&RUN assay^25^ to isolate live single CTCs from breast cancer patients and culture them individually. In one patient, two single CTCs cultured under regular media divided during the first two weeks: one cell divided once and the other divided twice (Figure 4). However, shortly after 2 weeks, cells started to die from both clones. In an attempt to rescue these clones, media was replaced with NAC containing media for one clone, which lead to cell proliferation for around 5 more weeks, forming a large colony of cells (Figure 4).

**Figure 4.**
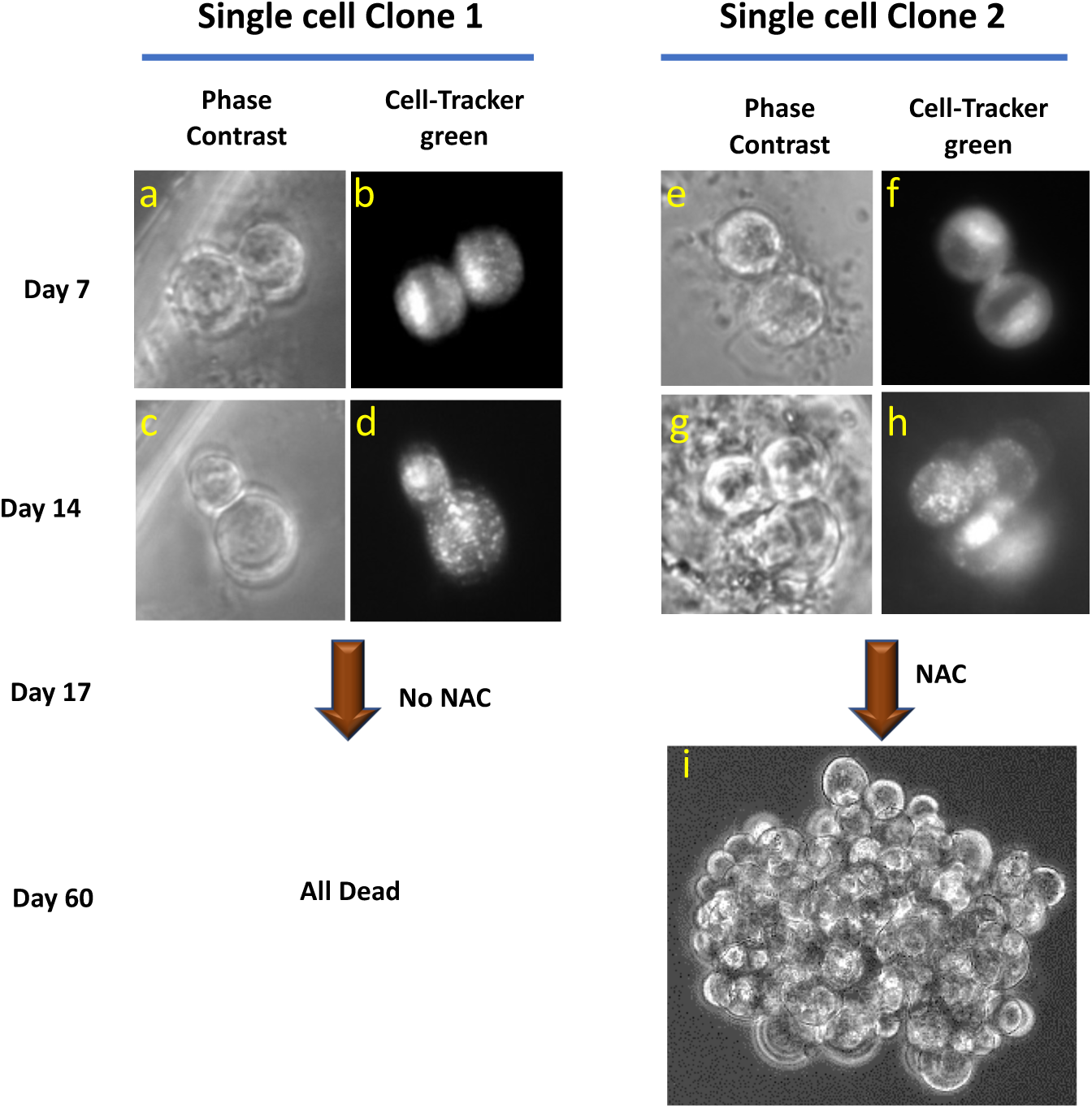
NAC treatment rescued a CTC clone isolated from a breast cancer patient. Two single CTCs were isolated from a tube of blood from a breast cancer patient were cultured in separate wells (single cell clone 1 (**a-d**) and single cell clone 2 (**e-i**)). Phase contrast microscope images for single cell clone 1 (**a and c**) and 2 (**e and g**) at day 7 and day 14 in culture under regular media. Images of Cell tracker green channel in single clone 1 (**b and d**) and 2 (**f and h**) at day 7 and day 14. **i)** phase contrast image for single cell clone 2 at day 60 (after 8 weeks of NAC treatment).

### Transcriptional analysis of growing and non-growing single cell clones

To identify the transcriptional changes influenced by these molecules, we performed RNA-seq analysis of pools of growing clones from control or small molecule treated conditions at days 6 and 13 (Supplementary Table S7). We analyzed 2 antioxidants (NAC and P4D8), 2 COX inhibitors (P1C2 and P1G7), and a combination of NAC and P1C2 in the BRx68 CTC line. Principal Component Analysis (PCA) showed a clear separation of conditions at day 13 versus 6 (Figure 5A). Interestingly, there is a dramatic heterogeneity in the control samples at day 13 compared to compound treated conditions. We identified differentially expressed genes (DEGs) at day 13 versus 6 across all conditions, and Ingenuity Pathway Analysis (IPA) showed enriched molecular and cellular function in metabolism, with lipid metabolism as the most significant category. Compared with control at day 6, NAC and P4D8 (2 antioxidants) treatment showed DEGs enriched in metabolic pathways including lipid metabolism, whereas COX inhibitors— P1C2 and P1G7—have DEGs related to cell cycle and cell movement (Figure 5B&C).

**Figure 5.**
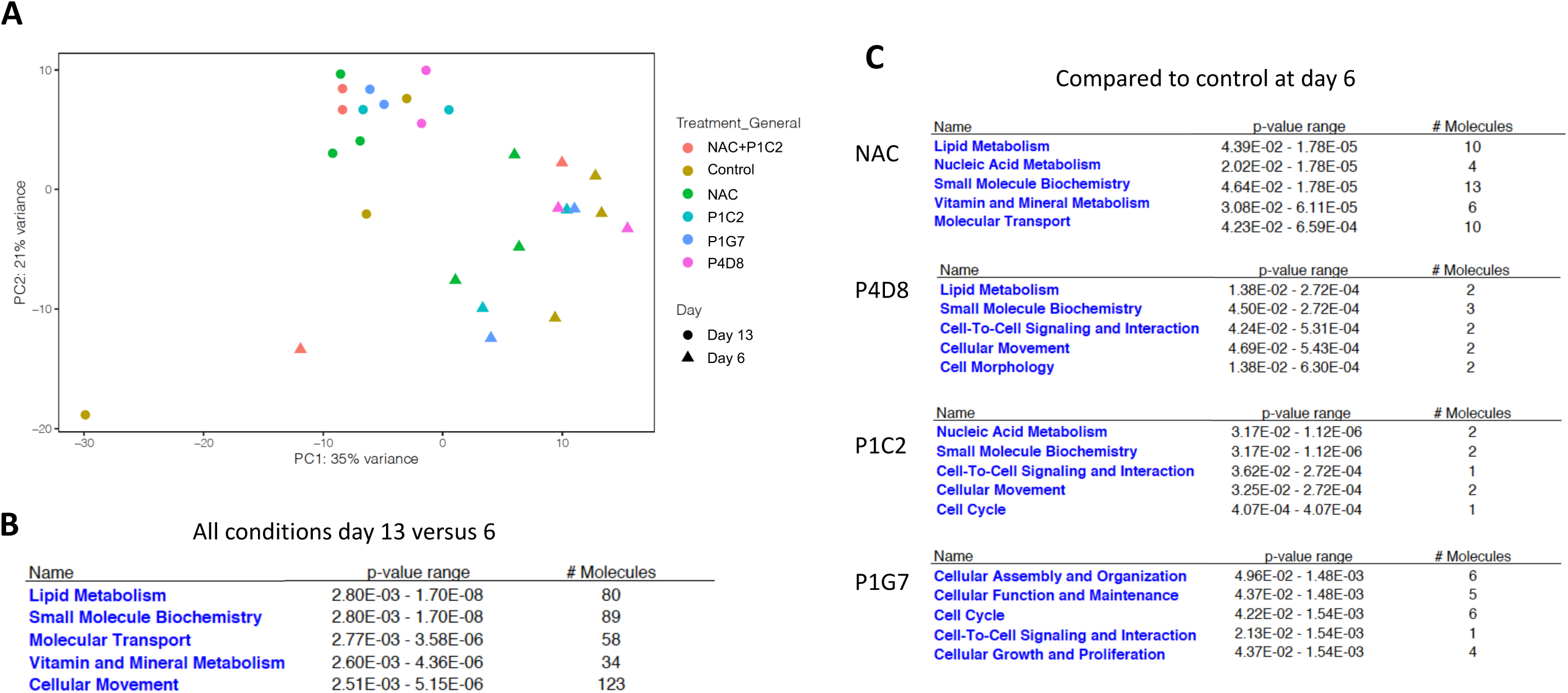
RNA-seq analysis of pools of clones at days 6 and 13. **(A)** PCA plot of RNA-seq results from pools of single cell clones at days 6 and 13 in control and molecule treated conditions, including 2 antioxidants (NAC and P4D8) and 2 COX inhibitors (P1C2 and P1G7). **(B)** Graphs of IPA analysis of enriched molecular and cellular functions of differentially expressed genes between days 13 and 6. **(C)** Graphs of IPA analysis of enriched molecular and cellular functions of differentially expressed genes between molecule treated conditions and control at day 6.

For a better understanding of how cell proliferation was induced by NAC since it showed more uniform effect across CTC lines, we also sequenced the non-growing cells from control and NAC. PCA analysis of the growing vs. non-growing cells showed a clear separation between both groups with high heterogeneity in the non-growing cells, which is expected as cell proliferation may have masked cell heterogeneity in the growing group (Figure 6A). The downregulated DEGs in non-growing cells compared with growing cells were enriched in cell cycle pathways (Figure 6B). IPA of DEGs in growing cells treated with NAC for 6 days compared to control non-growing cells showed enrichment of genes in stress and metabolism related pathways, albeit at low confidence of activation prediction (Figure 6C). Those pathways were not normally enriched in growing control cells when compared to non-growing control cells. Interestingly, at day 13, NAC treatment boosted the activation of cell cycle related pathways, much more prominent than that in control growing vs. non-growing cells (Figure 6D). This suggests that NAC helps cells alleviate the stress of being isolated individually at an early stage which is then followed with activation of cell proliferation.

**Figure 6.**
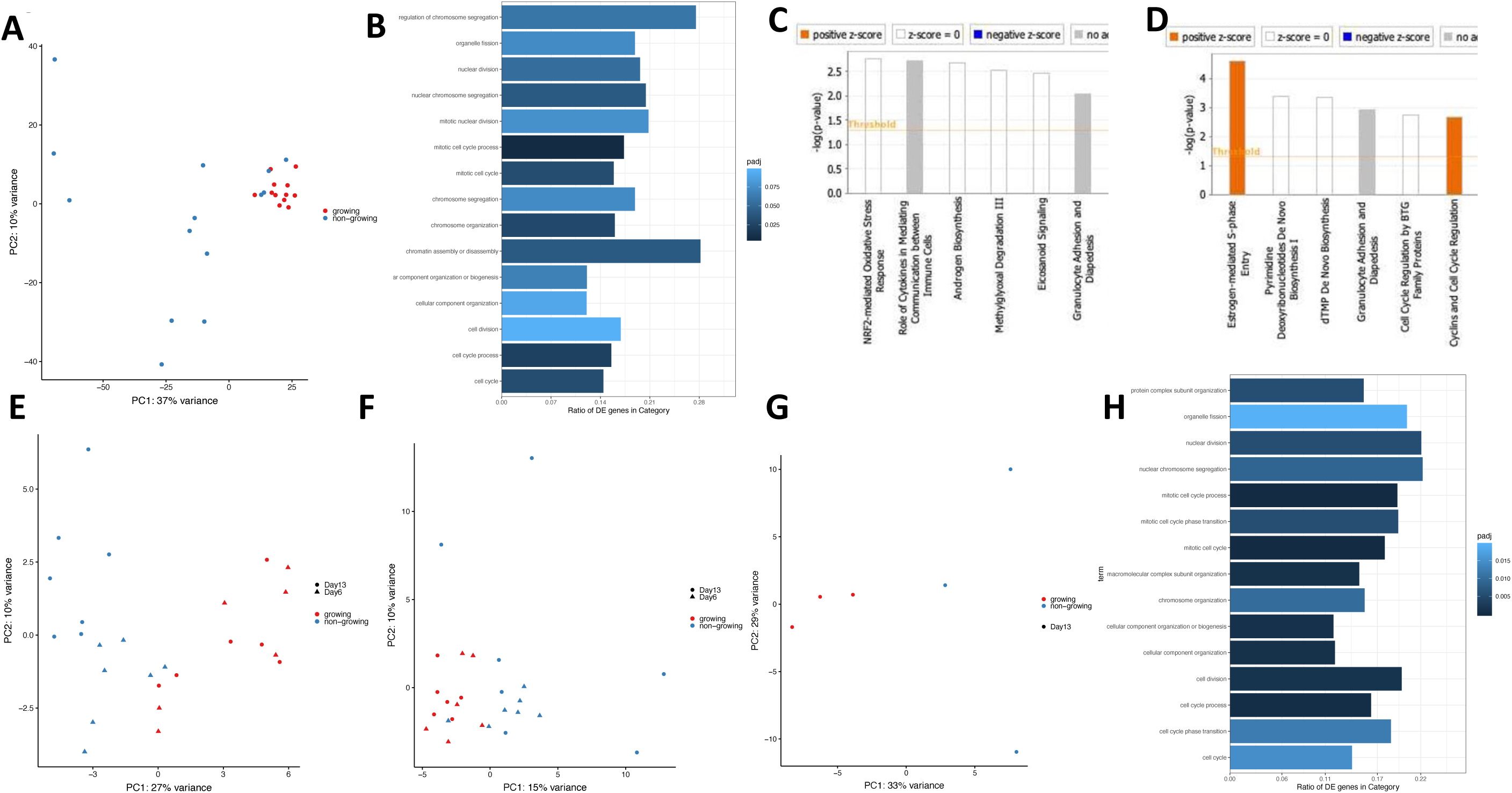
RNA-seq analysis of growing and non-growing cells. **(A)** Unsupervised PCA plot for both growing (red) and non-growing (blue) single cell clones. **(B)** Top pathways in Gene Ontology (GO) pathway analysis for downregulated DEGs in all non-growing cells compared with all growing cells. **(C-D)** IPA for DEGs in pools of growing and non-growing single cell clones treated with NAC for 6 days (**C**) or 13 days (**D**). **(E-G)** PCA plots for all samples based on previously published gene signatures of quiescent (**E**) and senescent (**F**), or for non-growing NAC treated cells at day 13 (**G**). **(H)** GO pathway analysis for downregulated DEGs in NAC-treated non-growing cells compared with growing cells.

Although NAC treatment significantly induced cell proliferation of single CTCs, many cells were not able to grow even under NAC treatment. This suggests that non-growing cells in both control and NAC groups may be quiescent, senescent or a combination of both. Indeed, PCA plots based on senescence and quiescent marker genes identified from a previously published study^26^ showed separation between growing and non-growing cells (Figure 6E and F), and more clearer when only comparing NAC treated cells at day 13 (Figure 6 G) with DEGs enriched more prominently in cell cycle pathways (Figure 6H), and 6 are among the 45 senescent markers (p = 0.079) (Supplementary Table S6). These analyses suggest that these non-growing cells contain a mix of quiescent and senescent cells and those did not grow after 13 days of NAC treatment are likely enriched for senescent cells.

## Discussion

In this study, we performed a small molecule library screening on single CTCs from the established breast cancer CTC lines to identify compounds that can promote single CTC expansion *in vitro*. We identified NAC at 300 μM concentration as the best compound for promoting single CTC growth from multiple patient-derived CTC lines. Given the rarity of CTCs isolated from patients, it is impossible to perform such large-scale screening studies directly from freshly isolated CTCs. Although our analysis used already established CTC lines and findings need to be further validated in additional patient samples, this is the first study using patient-derived CTCs to address a critical problem in expanding single CTCs *ex vivo*.

Our findings are consistent with the results of recent studies, reporting increased ROS levels in CTCs or cancer cells detached from matrix^12-14,17-20^. Other studies have demonstrated that this increase in ROS is in fact caused by detachment from the extracellular matrix or loss of cell-cell contact ^18,20^. The ROS increase needs to be mitigated via many mechanisms. For example, a fundamental change in citrate metabolism can promote proper redox balance for detached tumor cells^20^. Upregulation of free radical scavengers or exogenous antioxidants could also promote the survival of detached tumor cells in circulation^12,13,19^. In addition to oxidative stress, CTCs must also survive shear stress, immune attack, and physical constriction in the circulatory environment. Also, the isolation procedure, used before *ex vivo* expansion of CTCs, can be a source of stress. Therefore, antioxidant and free-radical scavengers could mitigate these stresses.

During single CTC expansion, RNA-seq analysis indicated significant metabolic changes. In small pools of growing clones of CTCs, there is also increasing transcriptomic heterogeneity, which can be reduced by molecules such as antioxidants or COX inhibitors. Antioxidants such as NAC induced changes in metabolism, including lipid metabolism, that may promote CTC proliferation. This is consistent with the previous report on the metabolic changes in tumor cells in suspension^13,20^. However, the exact role of genes involved in lipid metabolism needs further investigation in the context of single CTC *ex vivo* expansion. Moreover, RNA-seq analysis of NAC-treated cells also showed the presence of a heterogeneous population of single cells that are undergoing quiescence or senescence that could not be pushed into active cell cycle with NAC treatment alone. This presents a challenge for the effort in developing more efficient ways of culturing CTCs, and also raises the possibility that some of these cells—if survived the transit— may be able to lodge in secondary tissues in a relatively quiescent state.

We noted that the constraints on proliferation for those CTCs able to grow eventually are significantly reduced once single CTCs achieve a critical mass. This suggests that cell-cell contact mitigates stress, in line with recent findings on the survival advantage of CTC clusters^23,27^. This also indicates that compounds promoting CTC proliferation can be applied in the short term, and need not confound the downstream characterization of CTC biology. Our xenograft assay further confirmed this, given the similar tumorigenicity between several single cell clones from NAC and control groups.

The cell-type dependent effects of some compounds reflect the inter-patient heterogeneity of CTCs. While some compounds promote cell proliferation only in certain CTC lines, other compounds have either promoting or inhibitory effects, depending on the patient line. Previously, we have shown inter-patient heterogeneity in driver mutations in CTCs and associated patient-dependent drug susceptibilities^7^. While reducing oxidative stress is a common need for CTCs, other specific pathways are quite distinct among patients.

In summary, through a single cell screen using breast cancer patient-derived CTC lines, we identified NAC and other compounds with antioxidant properties that promote single CTC proliferation, likely by reducing the stress and altering cell metabolism to facilitate survival and growth. Although not applicable for every single CTCs, these compounds may be useful in improving the expansion of certain CTCs from a broader cancer patient cohort, thereby advancing our understanding of the biological properties of these rare and clinically important cells.

## Materials and Methods

### Cell culture

CTC lines were previously derived from metastatic breast cancer patients^7^. CTC lines were cultured in ultra-low attachment plates with RPMI 1640 medium, supplemented with EGF (20ng/ml), bFGF (20ng/ml), 1X B27 and 1X antibiotic/antimycotic, in 4% O_2_ and 5% CO_2_. Single CTCs were cultured in GravityTRAP™ ULA 96 well Plates (InSphero). Wells on the edges of the plates were not used to avoid influence from evaporation. Media with fresh compounds were exchanged every 3 days by inserting pipet tips onto the platform of the wells to prevent accidentally aspirating suspended CTCs at the bottom. CTC numbers were counted manually under an inverted microscope every 6 days.

### FACS sorting

Cells were pelleted and resuspended into single cell suspension in 1% BSA in PBS buffer with 7-AAD. Live single CTCs were sorted directly into 96-well GravityTRAP™ ULA Plates using aMoFlo cell sorter (Beckman Coulter). An inverted microscope was used to manually confim that there was 1 CTC per well, 1 hour after sorting.

### Compounds screening

Compounds were from the StemSelect library obtained from the Choi Family Therapeutic Screening Facility at the Eli and Edythe Broad CIRM Center for Regenerative Medicine and Stem Cell Research at USC. Each compound was given a code to ensure unbiased assessment and blinded to the investigators. Compound information is listed in Supplementary Table S1. In the first-round screening, compounds were used with 1μM concentration. Stocks of compounds (10mM concentration in DMSO) were stored in aliquots at −80^°^C. All compounds were thawed and refrozen for a maximum of 2 times. In the second-round validation, fresh compounds were dissolved in DMSO (Millipore sigma), and aliquots were stored at −20°C and used only once without refreezing. Only wells started from single cells on day 0 after sorting were used in the screening experiment. Data collection was done in batches.

### Spike-in experiment

Healthy volunteers’ blood samples were collected following protocols approved by the institutional review board (IRB) at the University of Southern California. GFP-positive CTC lines (50 cells/ml) were spiked into the blood samples from healthy volunteers, and RosetteSep™ CTC Enrichment Cocktail Containing Anti-CD56 was used to enrich CTCs from spiked-in samples. Isolated CTCs were cultured in GravityTRAP™ ULA Plate 96 wells. Each patient CTC cell line was processed by 2 different researchers.

### Single CTC clones

NAC clones were generated by treating single CTCs with 300μM NAC media for 24 days before switching to regular CTC media. Control single clone lines were established in the same batch using regular CTC media.

### Xenograft assay and Hematoxyline & Eosin staining

The animal protocol was approved by the Institutional Animal Care and Use Committee of the University of Southern California. Six-week old female NSG mice (Jackson Laboratory) were anesthetized with isoflurane and 20,000 GFP/luc-positive single clone cells in 100 μl of 1:1 PBS and Corning® Matrigel® Matrix (phenol-red free) were injected into the fourth mammary fat pad. To evaluate the growth of primary tumors, mice were intraperitoneally injected with 150 μl of d-Luciferin substrate at 30 mg/mL (Sid Labs), and imaged within 15 minutes. Signals from luciferase-tagged cells were monitored at day 0 after injection and weekly by *in vivo* imaging using IVIS Lumina II (Perkin Elmer) for 5 weeks. Mice were sacrificed after 8 weeks, and their organs were dissected and imaged. Primary tumors were collected and fixed with 10% formalin overnight and sectioned for 5um thickness. H&E staining was performed using Varistain Gemini ES Automated Slide Stainer in USC’s Histology Laboratory (HIST). Images were taken with a 20x objective in Keyence (BZ-II Analyser, Keyence).

### RNA-seq analysis

BRx68-GFP+ cells were sorted, with 1 cell per well, using the MoFlo cell sorter (Beckman Coulter). Single cells were cultured in the presence of CTC media containing either 1μM of P1C2, P4D8, or P1G7, or 300 μM of NAC or a combination of 1μM of P1C2 and 0.3 mM of NAC. Cells cultured in media containing DMSO served as a control. All cells were cultured for either 6 days or 13 days at 37°C, 5% CO2 and 4% O2, and media was changed every 3 days. For both day 6 and day 13 groups, clones with more than 3 cells at day 6 were harvested and pooled to a maximum of 50 cells as growing clones, and those less than 3 cells were collected as non-growing clones. Pooled samples were processed using SMARTer chemistry (SMART Seq® v4 Ultra® Low Input RNA Kit for Sequencing, Takara Clontech), according to manufacturer’s instructions to generate cDNA libraries for mRNA sequencing. All cDNA samples were run on a TapeStation system (High Sensitivity D5000 DNA Analysis Kit as per manufacturer’s protocol). cDNA libraries were prepared using the Nextera XT DNA Library Prep Kit (Illumina) with Nextera index kit index 1 (i7) and index 2 (i5) adapters. Libraries were sequenced on an Illumina NextSeq500 to obtain 75 bp-long single-end reads.

RNA-sequencing reads were trimmed for Nextera and Illumina adapter sequences using Trim Galore under default parameters. Trimmed reads were then mapped to the human genome build GRCh37 from Ensembl (ftp://ftp.ensembl.org/pub/grch37/current/fasta/homo_sapiens/dna/Homo_sapiens.GRCh37.dna_sm.primary_assembly.fa.gz) using STAR under optimized parameters for single-end sequenced data. Aligned reads were then counted via featureCounts (47) and piped into DESeq2 (48) for normalization to sequencing depth and downstream analysis. For purposes of producing the PCA plot, count data was transformed via the vst function to eliminate the experiment-wide trend of variance over mean and the plot was produced using ggplot2. For the PCA plot, batch effects were corrected using the function removeBatchEffect from limma. Differential expression analysis was performed controlling for batch effects. The contrast function was used to compare all conditions at day 13 versus day 6, each condition at day 13 versus day 6, or each treatment versus control at each time point. Genes with a False Discovery Rate (FDR) of 0.05 and log2 fold change of > 1.5 were piped into IPA for gene ontology analysis. Differentially expressed genes with fold change ≥ 2 and FDR ≤ 0.05 identified from a published study^26^ are used as marker genes for quiescent and senescence state.

### Statistical analysis

For the screening analysis, the number of cells grown over time per well per plate per batch was represented as AUC (area under the curve). A Wilcoxon-Rank Sum Test was performed to compare AUC of each drug to AUC of CT (control with CTC media only) or CD (control with matching amount of DMSO in CTC media) within each batch. P-values were adjusted by Benjamini-Hochberg Procedure to control the false discovery rate. In the validation experiments, drugs with adjusted p-values<=0.2 were considered as statistically different from CT/CD. Statistical tests were performed using R. For other experiments, data were analyzed with Student’s t test, and represent the means ± SEM of at least triplicate samples or averages ± SD of independent analyses, as indicated. P<0.05 was considered statistically significant. Statistical tests were performed with GraphPad Prism7 statistical software.

## Acknowledgments

We thank the USC Flow Core, USC Histology Core, USC Histology Laboratory and USC Biostatistics Core for their excellent technical support. We thank the USC Department of Animal Resources for caring of our experimental animals. We thank Jeffrey Boyd and Bernadette Masinsin from the USC Flow Cytometry Core for assisting in sorting single cells. We thank Mickey Huang from the USC Choi Family Therapeutic Screening Facility, Eli and Edythe Broad CIRM Center for Regenerative Medicine and Stem Cell Research at USC for helping to prepare all compounds used in the first round of screening. We are grateful to Cristy Lytal for editing our manuscript. This research was supported by the following: the National Institutes of Health (NIH) (DP2 CA206653 to M.Y.), the Stop Cancer Foundation (M.Y.), the PEW Charitable Trusts and the Alexander & Margaret Stewart Trust (M.Y.), the Wright Foundation pilot award (M.Y.), and the NIH grant T90DE021982 (trainee Y.A.). The project described was supported in part (core facilities) by the National Cancer Institute (grant P30CA014089).

## Author contributions

T.T. designed and performed most of the research and data analysis. O.I. and M.Y designed experiments. O.I. and G.L. optimized the conditions for the screening. M.K. performed, processed, and analyzed data for RNA-seq experiments. Y.A., A.T., and A.S. performed RNA-seq analysis. C.L. and R.S. performed statistical analysis. J.N. digitized and analyzed part of the raw data. M.Y. conceived and supervised the study. T.T., M.K. and M.Y. wrote the manuscript. All authors edited or commented on the manuscript.

## Data and materials availability

All data needed to evaluate the conclusions in the paper are present in the paper and/or the Supplementary Materials. RNA-seq data are deposited at GEO database (GSE134138). Additional data related to this paper may be requested from the authors.

## Figure legend

**Supplementary Figure S1.**
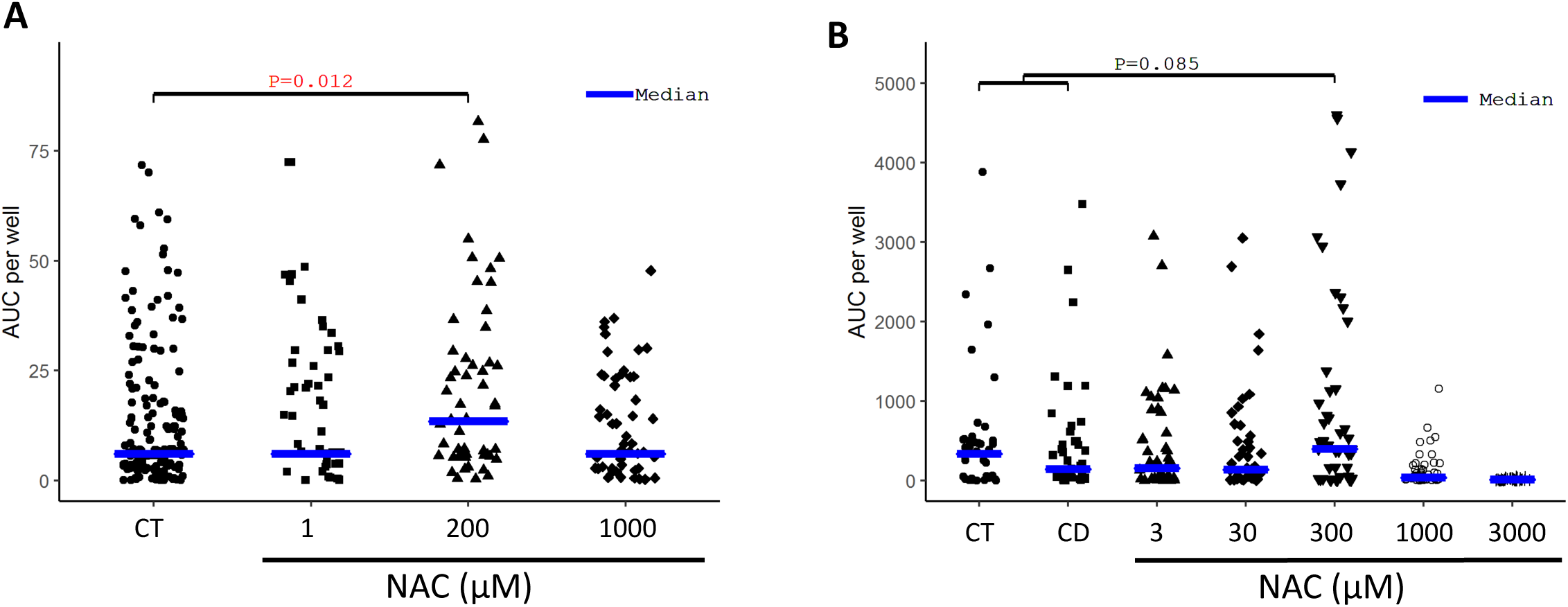
Optimization of NAC concentration. **(A**) Graph showing AUC measurement of the proliferation of single BRx68 cells over 24 days with various NAC concentrations. \**P*<0.05. **(B)** Graph showing AUC measurement of the proliferation of single BRx68 cells over 24 days with various NAC concentrations in a separate batch. \**P*<0.05.

**Supplementary Figure S2.**
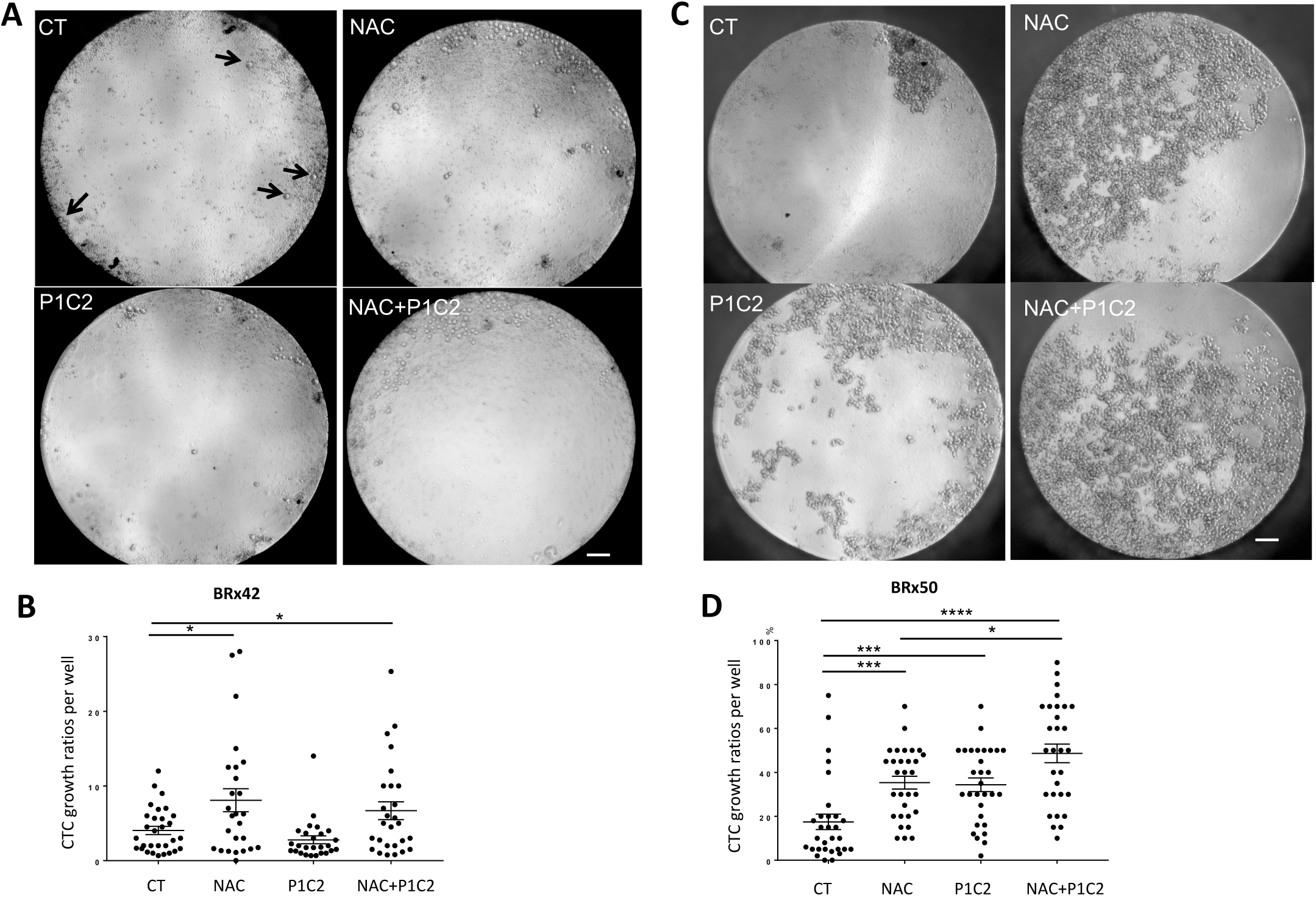
NAC and NAC+P1C2 combined promote growth of small numbers of CTCs isolated from blood samples. BRx42 **(A, B)** or BRx50 **(C, D)** spiked into healthy volunteers’ blood were isolated and cultured in either CTC media (control) or CTC media containing either NAC or NAC+P1C2 for 20 days. Representative images of BRx42 **(A)** or **BRx50 (C)** in 96 well plates at day 20. Graphs showing CTC growth ratio in each well of BRx42 **(B)** or BRx50 **(D)** at day 20. Each well contains 1–11 CTCs at day 0. Arrows point to single CTCs in the BRx42 CT condition. CT: N=30, P1C2: N=26, NAC: N=26 Mixture (P1C2+NAC): N=27. Scale bar: 200μm. mean ± s.e.m. *P* values were obtained with two-tailed unpaired t-test, F of F test both >0.05; ns, non-significant; * *P*<0.05; ** *P*<0.01; *** *P*<0.001; **** *P*<0.0001.

**Supplementary Table S1. Compound information**

**Supplementary Table S2. Compounds with better median AUC than control in first round**

**Supplementary Table S3. Results for the first round**

**Supplementary Table S4. Results for the second round**

**SupplementaryTable S5. AUC for all experiments**

**Supplementary Table S6. Overlap of senescence markers with DEGs of NAC day 13 non-growing and growing clones**

